# Image-guided alignment of consecutive multi-modal tissue slides

**DOI:** 10.64898/2025.12.02.690676

**Authors:** Benedetta Manzato, Claudio Novella Rausell, Gangqi Wang, Nina Ogrinc, Rosalie G.J. Rietjens, Marleen E. Jacobs, Christos Botos, Sebastien J. Dumas, Ton J. Rabelink, Ahmed Mahfouz

## Abstract

Multi-modal spatial data analysis often requires precise physical alignment of consecutive tissue sections, a process that can be challenging and typically relies on shared molecular markers or image recognition techniques. Here, we introduce COAST (Consecutive multi-Omics Alignment of Spatial Tissues), a method to reliably physically align consecutive tissue sections to produce a unified multi-modal molecular dataset suitable for downstream applications. COAST relies exclusively on the images associated with spatial data, eliminating the need for common molecular features or prior annotations. We demonstrate the effectiveness of COAST using spatial transcriptomics slides, where it achieves performance comparable to established uni-modal alignment tools. Applying COAST to spatial transcriptomics and metabolomics/lipidomics tissue sections from a mouse model of ischemia reperfusion injury allowed the investigation of lipid/metabolite features of transcriptionally-defined cell types. Overall, COAST offers a streamlined and integrative solution for multi-modal spatial data alignment.

## Introduction

Spatial omics revolutionized many areas of biology as it allows molecular profiling of cells within their native spatial context. This represents a significant advancement over single-cell omics, which, while highly sensitive and capable of identifying novel cell types through the analysis of dissociated cells, lack the critical spatial context ^1^. Among spatial omics technologies, spatially resolved transcriptomics (SRT) has emerged as the most widely adopted approach, similar to the prevalence of single-cell RNA sequencing (scRNA-seq) within the single-cell domain ^2^. However, recent advancements have extended spatial omics capabilities to other molecular layers, such as metabolomics and proteomics ^3,4^. Integrating these multiple omics enables the exploration of complementary molecular characteristics that contribute to cellular phenotypes, and is essential to understand their collective roles in driving complex biological phenomena ^5^.

Technologies which allow capturing multiple molecular layers simultaneously from the same tissue slide are rapidly developing ^6–8^. These approaches remain technically demanding and constrained by challenges such as differing slide preparation requirements between modalities and tissue degradation resulting in lower data quality ^9^. As a result, researchers often rely on applying different spatial modalities to adjacent or consecutive tissue sections.^10,11^. Aligning the consecutive sections is then essential to allow multi-modal downstream analysis. This task, however, is not trivial due to structural differences between sections as well as potential tissue deformation, distortion, and other artifacts introduced during the sectioning process ^12,13^.

Several methods have been proposed to address this problem of tissue alignment, which can be roughly divided into two categories. The first category achieves alignment by maximizing the similarity of the measured molecular features (e.g. gene expression in case of SRT) using graph neural networks ^14^, optimal transport ^15,16^, or gaussian processes ^17^ to align the feature distributions. These methods, however, are largely limited to aligning consecutive sections of the same spatial modality to construct virtual 3D tissue blocks ^18–20^ but do not generalize to multi-modal sections without matching features across sections (commonly referred to as diagonal integration ^21^). For example, CAST allows the integration of SRT and spatially resolved chromatin accessibility data by converting the accessibility data to gene activity scores, resulting in data loss. This approach is also not applicable to multimodal data in which molecular feature matching is more challenging (e.g. metabolomics and transcriptomics). Another class of methods identifies matching landmarks in tissue sections (i.e. locations with recognizable structures) that can be used to guide the alignment by applying affine transformation or geometric transformations ^22–24^. Tissue landmarks can be detected manually or automatically, which can be laborious and/or impractical when tissues lack clear visually-recognizable structures.

Here we present COAST (Consecutive multi-Omics Alignment of Spatial Tissues), a method to align consecutive sections profiled with any combination of multi-modal spatial omics technologies. COAST relies on the paired stained images generated alongside spatial omics data (i.e. nuclear staining or H&E) to construct a shared feature space in which spatial observations from separate sections can be aligned. Our approach reframes the alignment problem of multimodal spatial data from consecutive sections as a vertical integration task ^21^, where correspondence is established based on the similarity of matched image-derived features rather than unmatched molecular profiles. We benchmark COAST against existing uni-modal alignment methods using SRT datasets, highlighting its applicability to H&E and nuclear staining. To illustrate COAST’s true potential in multi-modal integration, we apply it to consecutive mouse kidney sections measured with Stereo-seq for spatial transcriptomics and MALDI-MSI for spatial metabolomics.

## Results

### Overview of COAST

COAST is a framework that takes consecutive spatial omics data and uses their paired staining images as an anchor to obtain a shared representation to map points of the two sections into a common coordinate framework (CCF), enabling the construction of unified uni- or multi-modal datasets of spatially aligned observations (Fig. 1a). From the two stained images, we start by extracting structural information that can serve as a shared representation. Vision Transformers have been shown to effectively capture cellular heterogeneity ^25^ and due to their ability to model both local and global context, they are well-suited for capturing long-range dependencies, thereby addressing morphological redundancy within tissue samples ^26^. Once we establish the shared image features for both datasets, we build on CAST, a method designed for aligning uni-modal consecutive sections, to map the coordinates of the query image/spatial data into a CCF. Specifically, we employ the module CAST Stack that performs an initial rigid transformation followed by a B-spline transformation based on embedding similarity. This process maps the coordinates of the query image/spatial data into a CCF. Then, using the transformed coordinates of the query image, spatial units are matched based on their physical proximity to observations in the reference image. The result is a unified uni- or multi-modal matrix, where features represent the combined set of measurements from both modalities, and the spatial observations are the matched locations from both sections following alignment.

**Fig 1:**
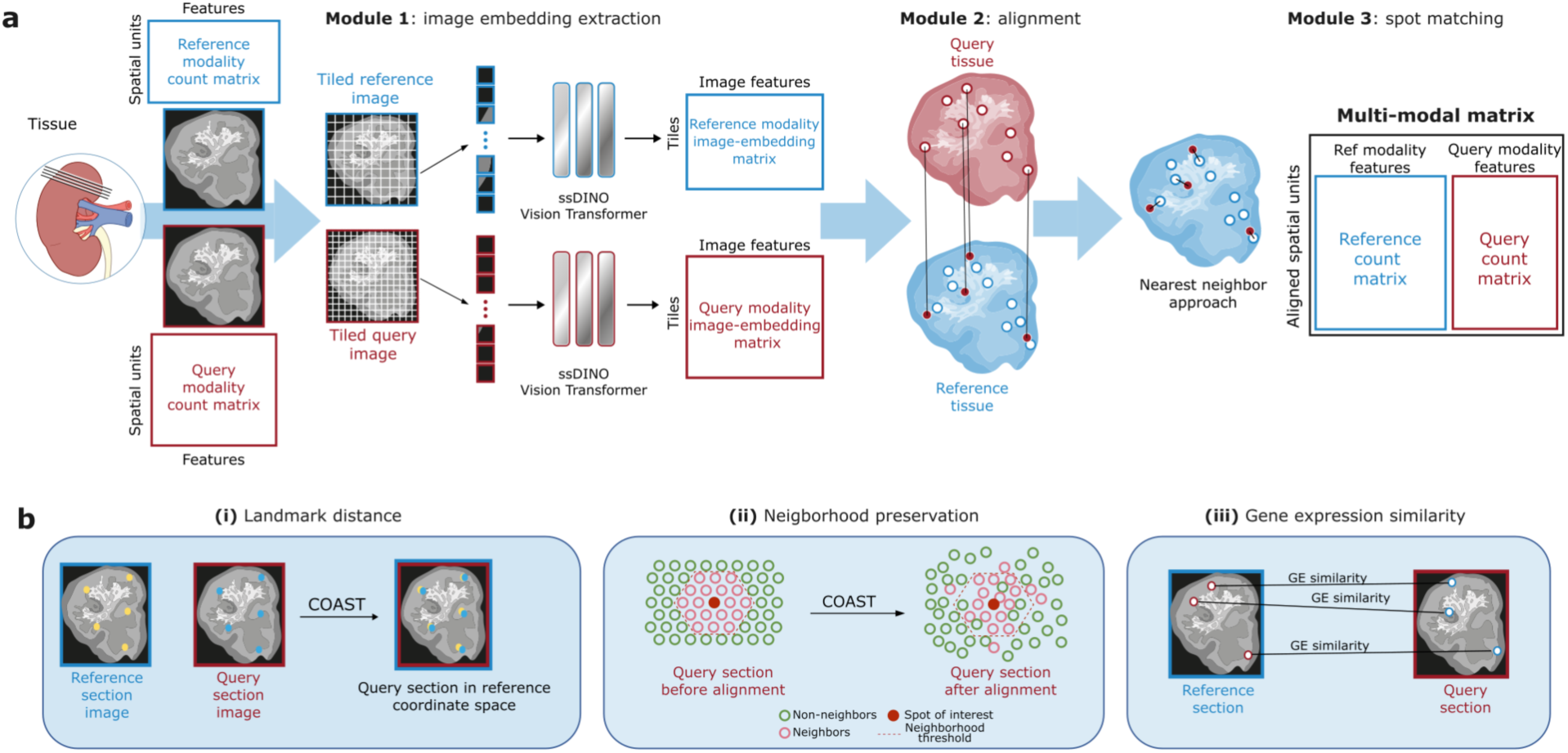
Schematic overview of Consecutive multi-Omics Alignment of Spatial Tissues (COAST). **a**. COAST takes as input two images that are tiled and fed separately to the ssDINO Vision Transformer to obtain an embedding of 384 image features for each image, where the size of the embedding is the size of the output layer of the vision transformer (Module 1). Query spots are mapped to the reference coordinate space based on the similarity between their image embeddings (Module 2). A unified uni- or mulit-modal matrix is created by matching query and reference spots based on their proximity (Module 3). **b**. The method is evaluated with three main quantitative metrics: (**i**) landmark distance, (**ii**) neighborhood preservation and (**iii**) gene expression (GE) similarity.

To quantify the quality of the resulting alignments, we measured the distance between landmark pairs post alignment. These landmarks served as a form of ground truth and were identified as corresponding points within the tissue that should overlap following alignment, such as tissue edges, tears, or recognizable anatomical structures visible from the staining. We then calculated the Euclidean distance between the corresponding reference and query landmarks after applying COAST, which should be minimal when the alignment is accurate (Fig. 1b(i)). To ensure that tissue integrity is maintained after alignment, we defined a neighborhood preservation score by comparing the third-order neighborhood of each query spot before and after alignment (Fig. 1b(ii)). Moreover, when aligning uni-modal data (e.g. both consecutive sections measure gene expression), we can assess the quality of the alignment based on the similarity of the molecular profiles between the matched spots (Fig. 1b(iii)). Importantly, we do not expect matched spots to have identical gene expression profiles, as a key characteristic of a good alignment is the preservation of (spatial) biological structure, not merely the optimization of gene expression or image feature similarity. This is especially important given that gene expression differences can naturally occur between tissue sections.

### COAST aligns 10X Visium consecutive sections based on the paired H&E images

Although COAST is designed to align multi-modal tissue sections, we first evaluated its performance using consecutive spatial transcriptomics sections to allow comparisons against existing uni-modal alignment methods. Specifically, we aligned three pairs of consecutive 10X Visium sections of the mouse brain: Sagittal Posterior, Coronal, and Sagittal Anterior (Fig. 2a, Supplementary Fig. 1). First, we assessed how well the image features generated by the transformer captures tissue structures by clustering the joint vision transformer embeddings. The resulting clusters capture some of the known tissue architecture, such as the cerebellar fold in both sections (Supplementary Fig. 2a,b). Next, we compared the performance of COAST to CAST and MOSCOT, two uni-modal alignment methods, as well as manual rigid alignment and the original unaligned sections. We selected a total of 19, 13 and 32 landmarks for the Mouse Brain Sagittal Posterior, Anterior and Coronal image sets, respectively (Supplementary Fig. 3a-f). The median distance between matching landmarks for COAST was 102.20, 12.39 and 135.29 pixels, which is relatively small considering the inter-spot distance for Visium is 69 (horizontally) and 138 (diagonally) pixels. For all the three datasets, COAST showed a significant improvement over the unaligned sections (Fig. 2b, Supplementary Fig. 1b) (d=-6.39, p < 0.001 for the Sagittal Posterior dataset; d=-5.73, p < 0.001 for the Sagittal Anterior dataset, d = -5.74, p < 0.001 for the Coronal dataset). Similarly, COAST outperformed rigid alignment in the Sagittal Posterior dataset (d=-0.60, not significant) and was significantly better in the Sagittal Anterior and Coronal dataset (d=-1.525, p=0.026 and d=-1.68, p=0.015). In all the datasets, COAST performed on par with CAST and with MOSCOT on the Sagittal Posterior dataset, while MOSCOT performed significantly worse than COAST (d=-2.61, p<0.001) and CAST (d=-2.92, p<0.001) in the Coronal dataset. Moreover, the number of spots matched between the consecutive sections is largely consistent across all the alignment tools, indicating that none of the methods heavily mismatched border regions during the alignment (Supplementary Fig. 5a,b). These results indicate that COAST can accurately align consecutive 10X Visium sections based on H&E images only, and achieves similar or better performance than CAST and MOSCOT without gene expression data.

**Figure 2:**
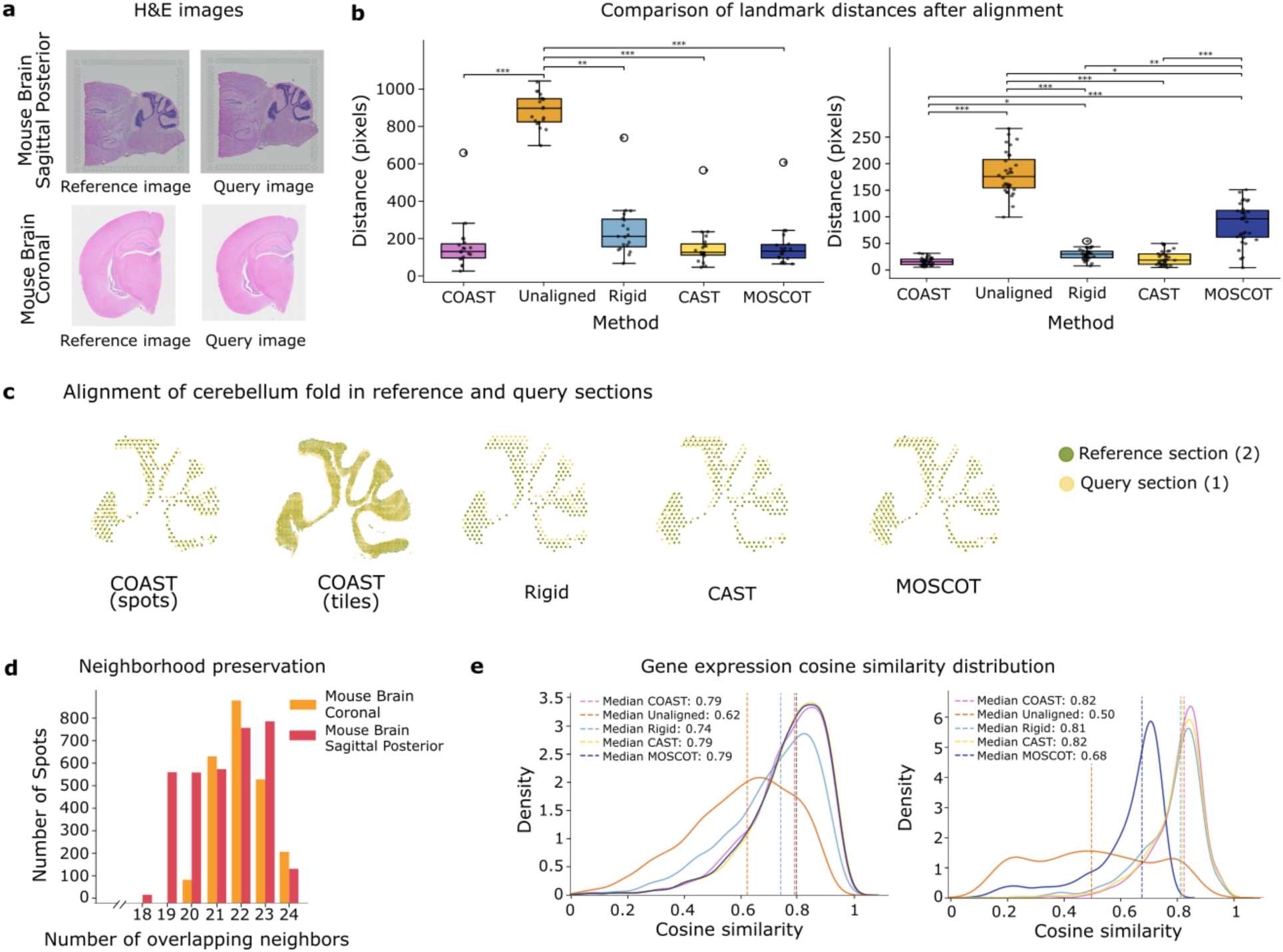
Aligning consecutive Visium slides using COAST. **a**: H&E images of the two consecutive mouse brain sections. Top: 10X Mouse Brain Sagittal Posterior, bottom: 10X Mouse Brain Coronal. **b**: Boxplot representing the distances between landmarks after registration via COAST, Rigid Alignment, Unaligned, CAST and MOSCOT. Left: Mouse Brain Sagittal Posterior, right: Mouse Brain Coronal. **c**: The cerebellar fold of Mouse Brain Sagittal Posterior dataset after alignment. **d**: Neighborhood preservation of the third order neighbourhood (24 spots) in the query section before and after alignment. **e**: Cosine similarity of gene expression of matched spots with COAST, rigid alignment, CAST and MOSCOT. Left: Mouse Brain Sagittal Posterior, right: Mouse Brain Coronal.

To further evaluate the accuracy of the alignment, we zoomed-in on the granular layer of the cerebellum. Since there are no anatomical annotations available, we first clustered the transcriptomic data of both sections from the Sagittal Posterior dataset and selected the cluster corresponding to the cerebellar granular layer (see Methods). We then plotted the location of the reference and query coordinates after alignment, for COAST both the spots and tiles alignment are shown for clarity (Fig. 2c). COAST, as well as CAST and MOSCOT, aligned the query and reference spots belonging to the cerebellar granular layer seamlessly, whereas there is a discrepancy when the spots are rigidly aligned. To assess neighborhood preservation we compared the third-order neighborhood of each query spot before and after alignment (Fig. 1b(ii)). For Visium data, the third order neighborhood corresponds to 24 spots. Across the three datasets, at least 75% and 83% of the original neighborhood (corresponding to 18, 18 and 20 out of 24 spots, respectively for Mouse Brain Sagittal Posterior, Anterior and Coronal) were preserved after alignment for every spot in the query tissue, demonstrating strong preservation of local tissue structure (Fig. 2d).

Since we are aligning uni-modal data we can assess the quality of the alignment based on gene expression similarity between the matched spots (as shown in Fig. 1b(iii)). Using this measure, in the Mouse Brain Sagittal Posterior dataset, COAST and the uni-modal methods CAST and MOSCOT outperform rigid alignment and unaligned baseline (Fig. 2e, left). For the Mouse Brain Coronal, COAST performs on par with CAST and rigid alignment while MOSCOT achieves lower performance as observed earlier through landmark evaluation (Fig. 2e, right). For the Mourse Brain Sagittal Anterior, COAST achieves better gene expression similarity than unaligned baseline and comparable similarity as the rigid alignment, CAST and MOSCOT (Supplementary Fig. 1e). Of note, unlike CAST and MOSCOT which utilize the expression data for alignment, and as such have an advantage when evaluating expression similarity of the aligned sections, COAST only uses the associated H&E images.

While the three Visium datasets shown in Figure 2 and Supplementary Figure 1 demonstrate good performance, these are relatively easy scenarios, characterized by well-preserved tissue morphology, minimal distortion, and consistent orientation across slides. To evaluate COAST under more challenging conditions, we artificially distorted the query images by applying a 90-degree rotation and a horizontal flip (around the x-axis), respectively. We then ran only the feature extraction and alignment modules on these altered images. COAST was still able to recover a meaningful alignment in all three cases, demonstrating its capacity to handle substantial spatial transformations (Supplementary Fig. 4). This suggests that the model captures underlying tissue structure in a way that is robust to such perturbations.

All together, these results show that COAST effectively aligns consecutive sections using the H&E stainings associated with SRT data, performing comparably to, and in some cases surpassing, existing unimodal methods incorporating molecular features.

### COAST generalizes to uni-modal data with nuclear staining images

The type of staining images accompanying spatial omics data varies across technologies. To further evaluate the generalizability of COAST to different stainings, we used the Mouse Embryo MOSTA spatial transcriptomics dataset. This dataset was generated using Stereo-seq ^18^, which acquires a nucleic acid staining images from each tissue section We applied COAST to two serial sections - E16.5_E2S6 and E16.5_E2S7 - sliced several hundred micrometers apart (Fig. 3a).

**Figure 3:**
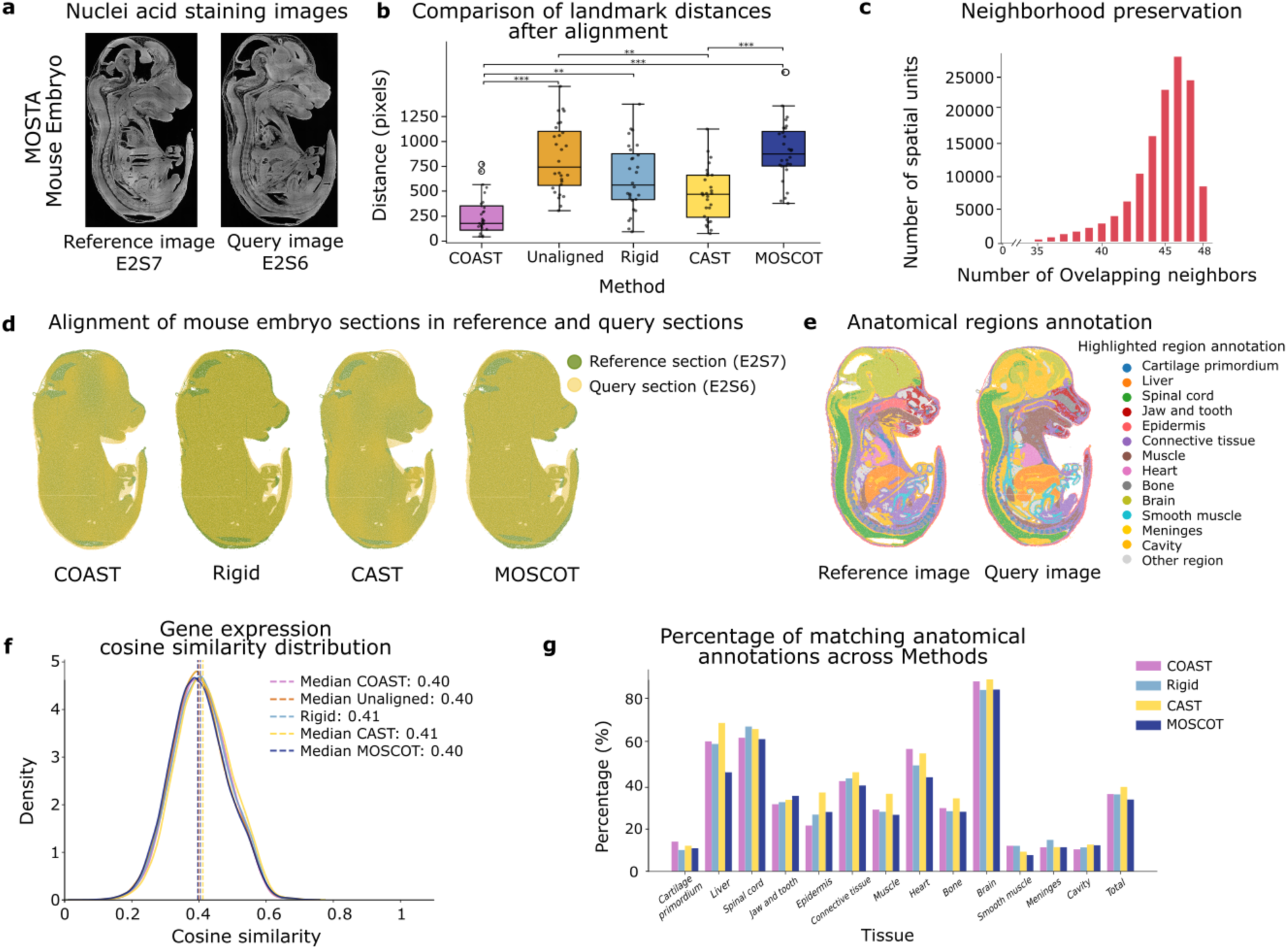
Aligning serial Stereo-seq slides using COAST. **a**: nuclei acid staining images of two serial mouse embryo sections. **b**: boxplot representing the distances between landmarks after registration via COAST, Rigid Alignment, Unaligned, CAST and MOSCOT. **c**: Neighborhood preservation of the third order neighbourhood (48 spots) in the query section before and after alignment. **d**: Spatial distribution of reference and query spots after alignment with COAST and benchmarking tools. **e**: Mouse Embryo sections colored by anatomical region (regions selected for downstream analysis only). **f**: Cosine similarity of gene expression of matched spots with COAST, rigid alignment, CAST and MOSCOT (raw counts, 3000 highly variable genes). **g**: Percentage of correctly matched anatomical regions after alignment with COAST and benchmarking tools.

We manually placed 28 landmarks and measured the pairwise distances between matching landmarks across sections after alignment (Supplementary Fig. 3g,h). COAST and CAST significantly outperformed MOSCOT with COAST also significantly outperforming rigid alignment (p<0.001, d>1.2) (Fig. 3b). Although the distances between matching landmarks were smaller with COAST compared to CAST, this difference was not significant. We also assessed neighborhood preservation to quantify tissue distortion. The spatial observations in Stereo-seq are organized in a grid-structure and therefore the third-order neighborhood includes 48 spots. We found that 100% of the spots retained at least 35 matching neighbors before and after alignment using COAST, indicating the method’s ability to uniformly reposition spots belonging to the query tissue (Fig. 3c). We calculated gene expression similarity between matched spots after alignment. All three methods as well as rigid alignment achieved a similar performance (Fig. 3f). The overall low similarity in gene expression compared to the 10X Visium data (Fig. 2e) is to be expected, given that the two sections are separated by several hundred micrometers. Additionally, the number of spots retained after alignment was comparable across methods, with the highest number retained by COAST (93.9%; 117,474 out of 125,090 bins) (Supplementary Fig. 5d).

Since the MOSTA dataset includes tissue annotations, we evaluated the quality of alignment by comparing the annotations of the matched spots. Since the distance between the sections is relatively large, there are significant changes in anatomical structures. We selected a subset of highly abundant annotations (more than 1000 cells) that were shared across both sections (Fig. 3e). This ensured that comparisons were based on robust and consistently represented populations. We then assessed the proportion of matched spots that retained the same region annotation post-alignment, presenting the results as a percentage of the total number of spots within each category (Fig. 3g). Overall, the percentage of matching annotations is not very high, similar to the overall low gene expression similarity noted above, due to the large differences between the sections. Importantly, there are no significant differences across the different alignment methods, with COAST consistently performing comparably to the other alignment methods.

These results demonstrate the generalizability of COAST to nuclei acid staining sections as well as its robustness to the distance between the consecutive sections, highlighting its applicability across spatial technologies and experimental settings.

### COAST aligns spatial transcriptomics, metabolomic and lipidomic data from the mouse kidney

To demonstrate COAST’s utility for studying complex biological processes through multimodal spatial data, we focused on kidney regeneration. Using an Ischemia-Reperfusion Injury (IRI) mouse model ^27^, we collected lipid and metabolite profile spectra via MALDI mass-spectrometry imaging (MALDI-MSI), and from a consecutive tissue section, 10 μm apart, we obtained whole-transcriptome gene expression data using Stereo-seq (Methods). The dataset includes three biological replicates. Both modalities are accompanied by corresponding DAPI and ssDNA-stained images (Fig. 4a).

**Figure 4:**
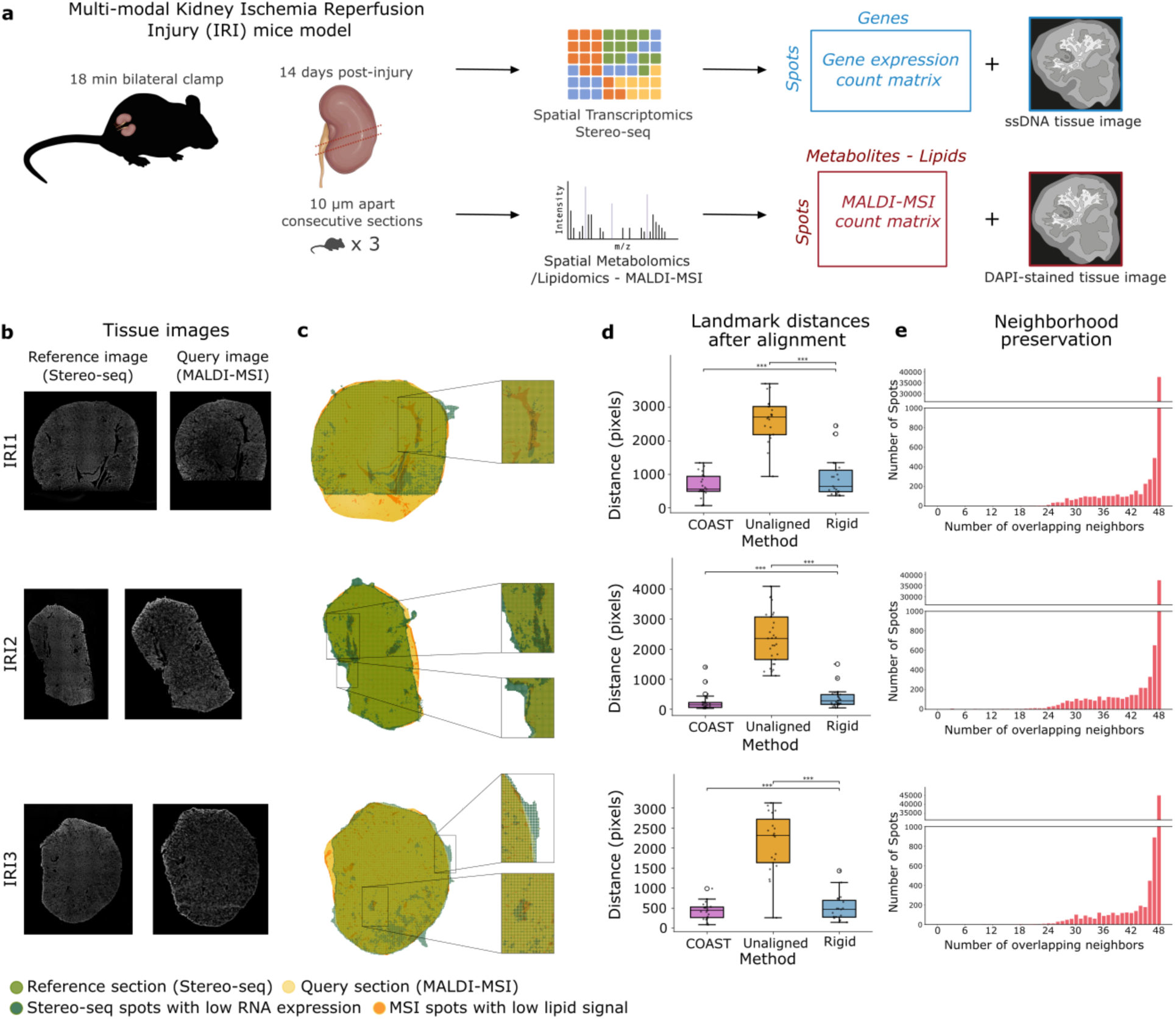
Aligning consecutive multi-modal slides using COAST. **a**: Multi Omics Ischemia Reperfusion Injury (IRI) experimental design. **b**: Tissue images. Left: ssDNA image of the reference tissues (Stereo-seq), right: DAPI-stained images of query tissues (MALDI-MSI). **c**: Alignment of the two consecutive sections with COAST (left) zoom-in of the alignment in selected regions of the tissues (right). **d**: Boxplot representing the distances between landmarks after registration via COAST, Unaligned, Rigid Alignment. **e**: Neighborhood preservation of the third order neighborhood (48 spots) in the query section before and after alignment.

For each pair of images (DAPI for MALDI-MSI and segmented-ssDNA for Stereo-seq, see Methods, Fig. 4b, Supplementary Fig. 6), we applied COAST to extract the image embeddings as described earlier (Fig.1a). We clustered the resulting embeddings for each image separately to assess whether they capture the same tissue structures. We observed that not all clusters were shared between the two modalities, likely due to the fact that, unlike previously analyzed image pairs, these images were acquired using different stainings, introducing a platform effect (Supplementary Fig. 2h). Nevertheless, when clustering was performed separately on the two consecutive section images, internal kidney structures remained clearly distinguishable (Supplementary Fig. 2i). Despite this platform effect, the alignment module of COAST was able to accurately match the two images across the three IRI kidney samples (Fig. 4c,d).

To evaluate alignment accuracy, we manually annotated corresponding anatomical landmarks across the three tissue pairs, identifying 21, 31, and 19 landmark pairs for IRI1, IRI2, and IRI3, respectively (Supplementary Fig. 2i,l). COAST consistently achieved significantly lower landmark distances compared to both unaligned and rigidly aligned sections, indicating more accurate spatial alignment (Fig. 4d).

To assess local structural preservation after alignment, we evaluated the overlap in spatial neighborhoods for each spot in the MALDI-MSI sections before and after alignment. For IRI1, 99.4% of spots retained at least 40 out of 48 of their original neighbors, and 95.6% retained all 48 neighbors. Similarly, in IRI2, 98.4% of spots retained at least 40 neighbors, with 93.5% preserving all 48. In IRI3, 98.7% of spots retained at least 40 neighbors, and 94.2% preserved all 48 (Fig. 4e). These results indicate that the alignment introduces minimal local distortion and preserves the original tissue architecture at a fine-grained spatial level across all samples.

These results demonstrate that COAST achieves high alignment accuracy while preserving tissue architecture, outperforming both unaligned baselines and, in some cases, manual rigid alignment. Importantly, these results demonstrate that COAST can align images even when input images originate from different nuclear stainings.

### COAST alignment identifies of multi-modal cell-type markers

Both transcriptomics and metabolomics have been used to investigate progression and repair mechanisms after IRI. These modalities were previously studied independently ^27–30^, but more recently efforts have shifted toward their integration, often across adjacent sections or without spatial context ^31–33^. Using COAST, we can now analyze these spatial multi-modal data simultaneously. Based on the resulting integrated data, we compared lipid and metabolite features and transcriptomic markers from matched spatial bins. This comparison was designed to highlight consistencies between the transcriptomic and metabolomic modalities, which should emerge if the alignment is sufficiently accurate at the bin level. For this task, lipid features were selected from the literature. ^27,34^ Lipid markers showed negative enrichment in regions annotated as tissue gaps (Fig. 5a), confirming the successful matching of gap regions across modalities. Among proximal tubule (PT) markers, the PT-S1/S2-specific lipid feature *m/z* 782.6 was enriched in PT-S1/S2, as well as in distal convoluted tubule (DCT) and connecting tubule (CNT) cells, consistent with their shared epithelial identity. Out of the three PT-S3 markers we included, two (*m/z* 772.6, 809.6) were enriched in PT-S3 cells, whereas *m/z* 462.4 was not. Injury-associated lipid features showed cell-type-specific enrichment. *m/z* 844.6 and 822.7 showed higher expression in injured tubules, even though no enrichment in the failed repair proximal tubules (FR-PT). Interestingly, *m/z* 822.7 and 838.7 are more expressed in PT-S3 compared to PT-S1/S2. This is consistent with the fact that the S3 segment operates under lower oxygen availability than S1/S2, rendering it more vulnerable to metabolic stress and injury ^35,36^. The thick ascending limb (TAL) shared multiple features (*m/z* 878.6, 744.6, 788.6, 750.6) with the collecting duct, in accordance with previous findings ^34^. Regarding the glomerular lipid features, *m/z* 556.4 (shared with Injured tubule and FR-PT) and 647.4 (shared with PT-S1/S2) show expression in the glomerular population, as previously shown in literature.

**Figure 5.**
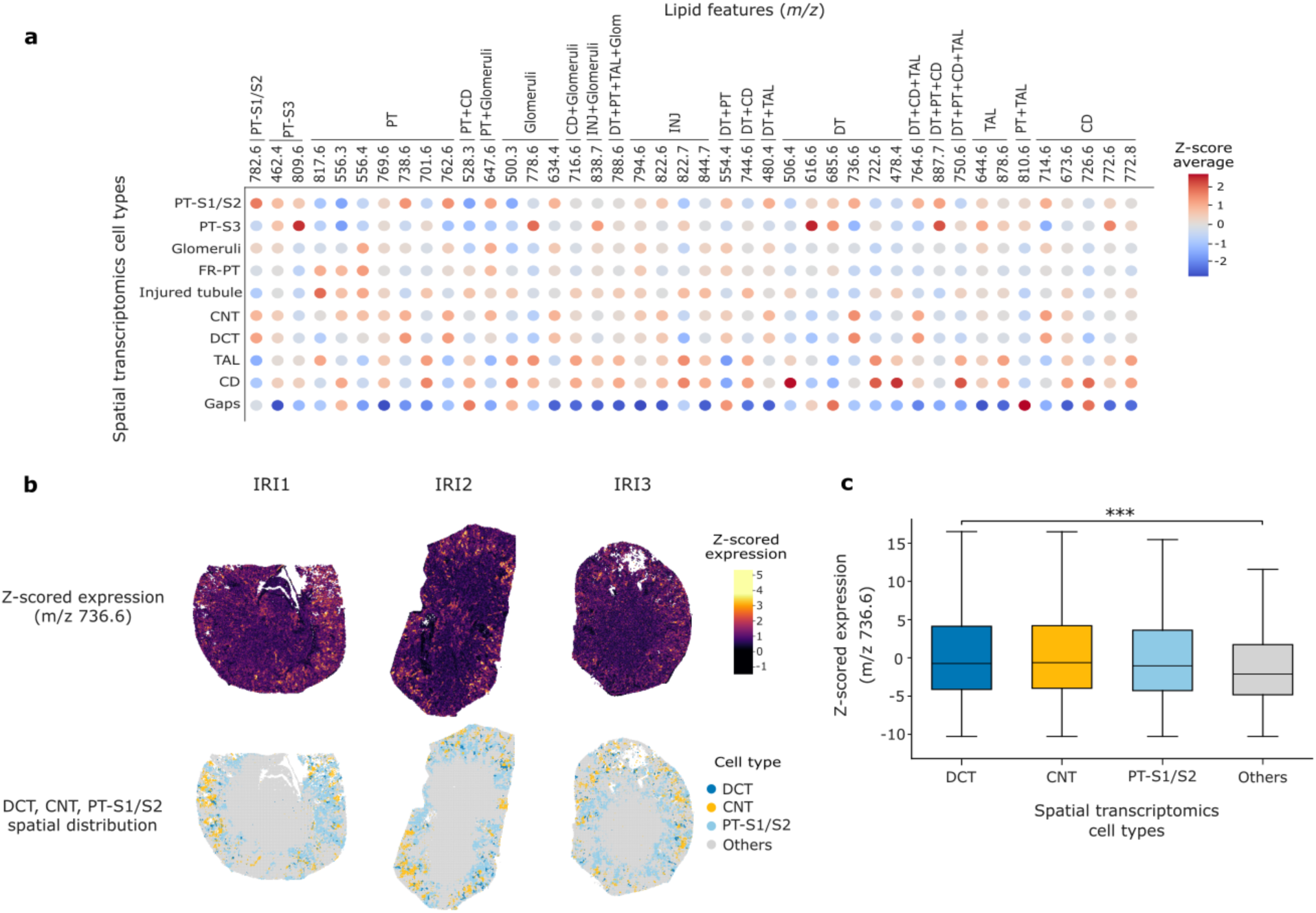
Downstream analysis on multi-modal dataset aligned with COAST. **a**: correspondence between lipid features (columns) and spatial transcriptomics-defined cell types (rows). Color indicates Z-score enrichment. The top column annotation indicates the cell type for which each lipid feature has been reported as a marker in the literature. **b**: Representative spatial maps showing the localization of lipid feature 736.6 (left) alongside corresponding transcriptomic cell type annotations (right). **c**: statistical analysis of lipid feature 736.6 in cell types of interest. Abbreviations: CD: collecting duct, PT-S1/S2: proximal tubule segment 1/2, PT-S3: proximal tubule segment 3,FR-PT: failed-repair proximal tubule, CNT: connecting tubule, DCT: distal convoluted tubule, TAL: thick ascending limb, INJ: injury.

Next we performed unbiased metabolite and lipid differential expression analysis between the transcriptionally-defined cell types (Supplementary Fig. S7). CNT and DCT exhibited nearly complete overlap in their identified markers, sharing 9 of the top 10 markers (m/z 185.0, 193.1, 195.1, 225.1, 239.2, 241.1, 267.1, 303.1, 539.2), supporting the close metabolic continuity of connecting and distal nephron segments. PT-S3 population displayed the most divergent lipidomic and metabolomic profile compared to other cell types, sharing none of the top 10 markers with any other cell type. Moreover, PT-S3 showed exclusive enrichment for multiple high-mass lipid species (m/z 809.6, 837.6, 838.7, 778.6, 888.8), highlighting a distinct lipidomic characterization. Injured tubules and FR-PT presented a mixed metabolic and lipidomic phenotype, with two overlapping markers (m/z 247.2, 435.2) and sharing several markers with CD, TAL, CNT, DCT and glomeruli (Supplementary Fig. S7).

Although only two features were unique to the glomerular population (m/z 554.355, 275.1), other non-unique features still showed enrichment (*m/z* 634.4, 556.4, 647.6). Importantly, also in this analysis, the gaps cluster showed no discriminant features, supporting its status as a region in the tissue with no lipid or metabolite expression.

Moreover, comparing the spatial distributions of lipid and transcriptomic markers showed an overall high concordance with cell type annotations obtained from transcriptomics. The PT-S3-specific lipid feature *m/z* 736.6 and transcriptomic DCT/CNT and PT-S1/S2 bins showed overlapping spatial distributions (Fig. 5b). Furthermore, the expression of m/z 736.6 was significantly higher in DCT compared to the other cell types (Fig. 5c). These results support the accuracy of alignment and highlight consistencies between lipid and transcriptomic measurements, while also revealing differences not detectable without spatial integration.

Overall, the downstream analyses carried out in this section using the COAST multi-modal output identified potential metabolite and lipid features that align known anatomical and functional structure of the kidney by reflecting biologically coherent patterns of feature sharing across related nephron sections.

## Discussion

We presented COAST, a method to align consecutive tissue sections using only image data. In contrast to existing approaches, COAST does not rely on matched molecular profiles or cell-type annotations, avoiding potential biases and expanding applicability to diverse multi-modal datasets. Despite using only image features for alignment, COAST effectively aligns spatial spots with similar visual characteristics and, notably, achieves high similarity in gene expression between matched spots in the case of uni-modal datasets. It also preserves tissue architecture and minimizes unnecessary spot displacement, as demonstrated by the neighborhood preservation metric. COAST is applicable to different staining types, making it applicable to diverse spatial technologies. This generalizability indicates that image data and the use of pre-trained ssDINO vision transformer can reflect meaningful internal tissue structure for guiding the alignment. Across three quantitative metrics - landmark distance, gene expression similarity and neighborhood preservation - as well as qualitative visualizations, we showed that COAST performs on par with established unimodal tools and surpasses manual rigid alignment, which is often time-consuming and less precise. Finally, we illustrated how COAST output can be used for downstream multimodal analyses, revealing lipids and metabolites differentially expressed that reflect the nephron structure, such as a strong overlap between CNT and DCT, and recapitulate known injury-associated features, including the enrichment of the injury markers *m/z* 822.7 and 838.7 in PT-S3 compared to PT-S1/S2, consistent with the ischemia reperfusion injury phenotype reported in literature.

COAST offers a time- and cost-efficient solution that, while requiring GPU resources, runs rapidly. Its modular architecture, comprising three main components, enables flexibility: individual modules (such as the vision transformer or the spot-matching strategy) can be adapted or replaced depending on the technology or study objective.

One current limitation is that the method benefits greatly from well-defined tissue borders; as a result, performance is reduced when aligning smaller tissue regions or immunofluorescence samples where staining is stronger in the tissue core than at the edges. Moreover, while a satisfactory alignment was ultimately achieved for the multi-modal dataset, it required image preprocessing steps to mitigate platform effects arising from differences between imaging systems.

Finally, establishing a reliable physical alignment between consecutive tissue sections is a critical prerequisite for multi-modal downstream analyses and for drawing biologically meaningful conclusions. COAST fulfills this need by providing accurate physical alignment that preserves tissue architecture and spatial context across modalities, constituting a robust, flexible and streamlined pipeline that performs strongly without relying on prior knowledge or molecular annotations.

## Methods

### COAST algorithm

COAST takes as an input the high-resolution images associated with the spatial data and the spatial count matrices, where the x and y coordinates are present in the metadata and match the coordinates of the image. The method includes three main modules:

#### Feature extraction

Initially, both images are divided into tiles, that are obtained by taking a square of pixels around each spatial unit centroid, taken from the x and y columns in the metadata, as well as a grid-like manner across the whole image, including the background (“in-tissue” and “grid” tiles, respectively). *Tile_size* parameter controls the size of the tile in pixels and should reflect the capture area (in pixels) of the spatial units in the dataset (i.e. the pixel gap between their centroids). *Step_size* parameter controls how much the sliding window moves between consecutive tiles in the grid-search; it should be chosen to take a number of tiles from the background proportional to the overall image resolution and should be large enough to avoid generating unnecessary tiles and slow down computations. More in detail, we took tiles of size 80x80 and step size 120 for Mouse Brain Sagittal Posterior and Mouse Brain Sagittal Anterior, given that the gap between spots for Visium is 28 (horizontally) and 120 (vertically), in pixels. We used tile size 16x16 and step size 32, considering the gap is 10 (horizontally) and 18 (vertically). The MOSTA Mouse Embryo dataset achieves cellular resolution (with an average gap of only ∼1.03 pixels between spatial units). Considering also the high resolution of the associated images, we used tile size (32x32) large enough to still capture characteristics from the immediate surrounding of each cell, and step size 224 to keep the total number of tiles at a manageable level for downstream computations. Using the same criteria, for the multimodal IRI datasets, we took tiles size 40x40 and step size 80 for Stereo-seq and tile size 8x8 and step size 80 for MALDI-MSI sections, being the bin distance 40 for Stereo-seq and 7.8 pixels for MALDI-MSI.

Tiles are resized to 224x224 to be compatible with the input of the vision transformer and image features are subsequently extracted using an ImageNet-pre-trained Vision Transformer (ssDINO) ^37,38^ resulting in a dataset with 384 image features for each tile. The method has an optional argument to add a 300 pixels padding to one side of the image to help the alignment. Padding has been added on the left side to the three multi-modal IRI datasets.

#### Alignment

Building on CAST, a method for alignment of uni-modal consecutive sections, we use the module CAST Stack to map the coordinates of the query section in the reference coordinate system based on the similarity of the 384-dimensional transformer embeddings. First, a gradient-descent-based rigid alignment (translation, rotation, scaling and reflection) is applied to have both sections in the same scale and coordinate system, followed by a free-form deformation (FFD), a robust non-linear warping approach, to handle local morphological differences among tissue samples. Instead of implementing the alignment by satisfying every cell at its optimum, CAST Stack prioritizes preserving biologically meaningful tissue structure, through the optimization of molecular data. In our case, we replace the molecular data in this step with the image embeddings obtained from the vision transformer.

Due to the large number of tiles in high-resolution images we use the function CAST.sub_data_extract to select a random subset of tiles (20,000) and CAST Stack is applied on the dataset at a reduced size. After generating the transformation parameters, we then apply the calculated alignment transformation to all cells in the original sample at the full resolution.

#### Spot matching

After alignment, only the tiles centered around molecular data (“in-tissue” tiles) are retained by filtering out tiles originally obtained through grid-like search (“grid” tiles). Each spot in the reference section was matched to its closest spatial unit in the query section by querying the nearest neighbor in the *transformed* spatial coordinates, using a k-d tree data structure for efficient Euclidean distance search. A match was retained only if the closest spot fell within a defined maximum distance threshold; spots without a valid match within this threshold were excluded from further multimodal integration. This maximum distance (max_distance) serves as a key hyperparameter and should be chosen to reflect the physical distance, measured in image pixels, between the centroids of adjacent spatial units, based on the resolution and layout of the specific spatial transcriptomics platform. For example, we used a threshold of 90 pixels for the three 10X Visium datasets, 40 pixels for MOSTA Mouse Embryo and 40 pixels for the multi-modal IRI dataset.

If a query spot maps to more than one reference spot or if the query section has a higher spatial resolution than the reference, this causes the reference spots to be duplicated, resulting in an “artificial upscaling” of the reference spatial units. The final output is a set of matched spot pairs across modalities, which enables the creation of a multimodal (or combined uni-modal) feature matrix combining molecular information across aligned spatial locations (Fig. 1c).

#### Clustering on image features and visualization

Following the feature extraction module, we performed unsupervised clustering on the image-derived features from each pair of tissue sections. The feature vectors from both sections were concatenated into a single matrix and standardized using z-score normalization. K-means clustering was then applied to the combined dataset. Using the associated spatial coordinates, we visualized the resulting clusters by assigning each a unique color and displaying them as scatterplots.

#### Implementation

COAST was implemented in Python 3.10.14 and developed to run on a CUDA-enabled GPU. Key libraries used included PyTorch for model loading ^39^, Hugging Face for the DINO Vision Transformer backbone, CAST for the alignment step ^14^, torchvision and PIL for image handling and numpy and pandas for data manipulation. More information can be found on the COAST github page.

COAST is scalable and applicable to a wide range of dataset resolutions. Table 2 reports the runtime for all the COAST modules (feature extraction, CAST alignment, spot matching and aligned dataset saving and results visualization) on all datasets used in this study. With GPU, COAST runs within 6 minutes for Visium datasets and around one hour for cellular resolution dataset MOSTA Mouse Embryo and for the multi-modal high-resolution IRI dataset.

### Datasets

#### 10X Genomics Mouse Brain Sagittal Posterior dataset

This data was taken from the 10X website (see Data Availability). In detail, fresh frozen adult mouse brain tissue sections were processed using the 10X Genomics Visium Gene Expression protocol (Spatial 3’ v1). The tissue was sectioned in the sagittal orientation (posterior region). Two consecutive sections were used: section 1 with 3,355 in-tissue spots and section 2 with 3,289 in-tissue spots. Gene expression was quantified for 32,285 genes. The associated H&E images have a resolution of 11,620 × 11,607 pixels for both sections.

The dataset was preprocessed using Scanpy ^40^. Cells with fewer than 5,000 total counts or more than 35,000 total counts were removed, and genes expressed in fewer than 10 cells were excluded from downstream analysis. Following filtering (by total counts and genes by detection rate), normalization and log-transformation, we performed PCA, neighbor graph construction, UMAP embedding, and unsupervised clustering separately on the two sections using the Leiden algorithm (default resolution 1). Spatial visualization of the resulting clusters was conducted using high-resolution histology images overlaid with spatial coordinates. The cluster overlapping the cerebellum fold was manually identified and retained for further analysis. This cluster contained 197 spots in section 1 and 201 spots in section 2. To generate the plot in Figure 2c(ii), we identified the tiles corresponding to the cerebellar fold cluster, as determined from clustering the image-derived features (Supplementary Fig. 3b, cluster 1).

#### 10X Genomics Mouse Brain Coronal dataset

This data was taken from the 10X website (see Data Availability). In detail, fresh frozen adult mouse brain tissue sections were processed using the 10X Genomics Visium CytAssist Gene Expression protocol (Visium V4 Slide - FFPE v2). The tissue was sectioned in the coronal orientation. Two consecutive sections were used: section 1 with 2,310 in-tissue spots and section 2 with 2,235 in-tissue spots. Gene expression was quantified for 32,285 genes. The associated histology images have a resolution of 1,873 x 2,000 pixels and 1,692 x 2,000 pixels, respectively for sections 1 and 2.

#### 10X Genomics Mouse Brain Sagittal Anterior dataset

This data was taken from the 10X website (see Data Availability). In detail, fresh frozen adult mouse brain tissue sections were processed using the 10X Genomics Visium Gene Expression protocol. The tissue was sectioned in the sagittal orientation (anterior region). Two consecutive sections were used: section 1 with 2,695 in-tissue spots and section 2 with 2,823 in-tissue spots. Gene expression was quantified for 32,285 genes. The associated H&E images have a resolution of 11,620 x 11,607 pixels for both sections. Preprocessing was performed in the same way as 10X Genomics Mouse Brain - Sagittal Posterior.

#### MOSTA Mouse Embryo (Stereo-seq Platform)

We analyzed the E16.5 embryonic mouse dataset, previously published by Chen et al. ^18^ The data were acquired using the Stereo-seq platform on frozen consecutive sections spaced several hundred micrometers apart, each 10 μm thick. We used sections E16.5_E2S6 (125,090 spots) and E16.5_E2S7 (128,143 spots), as they are the only consecutive sections with associated nuclei acid staining images available. Both sections measured gene expression across 27,574 genes. The authors of the paper normalized and log-transformed the raw count matrix using Scanpy ^40^ (sc.pp.normalize_total and sc.pp.log1p). Moreover, anatomical region annotations were available and were used for Figure 3g (Table of markers available on MOSTA dataset website). The DAPI-stained images had dimensions of E2S6: 16,385 x 27,200 pixels and E2S7: 16,504 x 28,574 pixels. Since the Stereo-seq data were provided in bin-50 resolution, we reprocessed the spot coordinates to align them with pixel-based image coordinates, using the Bin1 matrix files available on the dataset website (E16.5_E2S6_GEM_bin1.tsv.gz, E16.5_E2S7_GEM_bin1.tsv.gz)

### Benchmark methods

#### Rigid Alignment

We performed rigid alignment in Python by manually translating, rotating and/or scaling the x and y coordinates of the spatial units of the query section to obtain an optimal visual alignment to the coordinates of the reference section (Table 1).

**Table 1:** Types of transformations and their application order (indicated in parentheses) applied to the query coordinates for rigid alignment.

|  | Query section | X-axis translation (in pixels) | Y-axis translation (in pixels) | Rotation (degrees) | Scaling |
| --- | --- | --- | --- | --- | --- |
| Mouse Brain Sagittal Posterior | 2 | + 100 | + 590 | - | - |
| Mouse Brain Coronal | 2 | - | - | - | Rescale x and y on section 2's bounds |
| Mouse Brain Sagittal Anterior | 2 | + 110 | - 730 | - | - |
| MOSTA Mouse Embryo | E16.6_E2S6 | - | + 200 | - | - |
| IRI1 | MALDI- MSI | +10000 (1)<br>+ 300 (5) | +18000 (2)<br>+400 (6) | 10° counter-clockwise (3) | x stretched by 2.5, y stretched by 2.3 (4) |
| IRI2 | MALDI- MSI | +15000 (1) | + 8000 (2) | 15° counter-clockwise (3) | around centroid, both x and y (4) |
| IRI3 | MALDI- MSI | + 5500 (1) | + 8000 (2) | 20° counter-clockwise (3) | around centroid, both x and y (4) |

**Table 2:** Runtime, number of spatial units and image size for the datasets used in the study.

| Dataset | Number of spatial units | Image size (in pixels) | Number of tiles | Runtime |
| --- | --- | --- | --- | --- |
| **10X Mouse Brain Sagittal Posterior** | S1: 3,355<br>S2: 3,289 | S1: 11,620x11,607<br>S2: 11,620x11,607 | S1: 12,764<br>S2: 12,698 | 5.06 min |
| **10X Mouse Brain Coronal** | S1: 2,310<br>S2: 2,235 | S1: 1,872x2,000<br>S2: 1,692x2,000 | S1: 6,072<br>S2: 5,574 | 4.05 min |
| **10X Mouse Brain Sagittal Anterior** | S1: 2,695<br>S2: 2,823 | S1: 11,620x11,607<br>S2: 11,620x11,607 | S1: 12,104<br>S2: 12,232 | 6.11 min |
| **MOSTA Mouse Embryo** | E2S6: 125,090<br>E2S7: 128,143 | E2S6: 16,609x27,200<br>E2S7: 16,728x28,574 | E2S6: 134,118<br>E2S7: 137,615 | 65.24 min |
| **Multimodal dataset (IRI1)** | Stereo-seq: 48,347<br>MALDI-MSI: 240,255 | Stereo-seq: 12,659x11,336<br>MALDI-MSI: 4,352x5,120 | Stereo-seq: 70,925<br>MALDI-MSI: 243,775 | 67.12 min |
| **Multimodal dataset (IRI2)** | Stereo-seq: 40,918<br>MALDI-MSI: 161,596 | Stereo-seq: 9,710x15,063<br>MALDI-MSI: 4,352x4,864 | Stereo-seq: 51,124<br>MALDI-MSI: 164,951 | 72.34 min |
| **Multimodal dataset (IRI3)** | Stereo-seq: 40,494<br>MALDI-MSI: 165,870 | Stereo-seq: 10,444x12,561<br>MALDI-MSI: 3,840x4,352 | Stereo-seq:<br>MALDI-MSI: | 65.56 min |

#### MOSCOT

We ran MOSCOT with default parameters. For all spatial transcriptomics datasets, alignments were performed using both affine and warp transformations separately and the alignment yielding the most accurate visual concordance was selected for downstream analysis. Final coordinates for the Mouse Brain Sagittal Posterior, Mouse Brain Sagittal Anterior, and MOSTA datasets were derived from affine transformations. For the Mouse Brain Coronal dataset, the final coordinates were obtained using the warp transformation.

#### CAST

CAST Mark and Stack were run on the uni-modal benchmarking datasets with the default parameters. The MOSTA Mouse Embryo dataset has been downsized with the function CAST.sub_data_extract (20,000 spots).

For all benchmarked methods, spatial units were matched between the query and reference in the same manner as the spot matching procedure described in the COAST algorithm section (see Methods). For the comparison to unaligned sections, given 10X Visium Mouse Posterior and Anterior, MOSTA Mouse Embryo image sets were in the same coordinate scale (i.e. same tissue coordinate magnitude), no transformation was applied. The two consecutive IRI kidneys are in completely different coordinate spaces therefore a center of mass transformation was applied to the coordinate of the MSI tissues in order to overlap the Stereo-seq tissues to some extent.

### Evaluation metrics

#### Landmark Alignment Accuracy

To evaluate the accuracy of spatial alignment across different methods, we measured the Euclidean distances between manually annotated anatomical landmarks and their corresponding points after alignment. For each alignment method, we first assessed the normality of the distance distributions using the Shapiro-Wilk test. Depending on the outcome, we selected either a one-way ANOVA (for normally distributed data) or a non-parametric Kruskal-Wallis test (for non-normal data) to compare alignment performance across groups. A significance level of 0.05 was used throughout.

Following a significant result in the group-level test, we performed pairwise comparisons using Dunn’s test with Bonferroni correction to adjust for multiple testing. We computed Cohen’s d for all pairwise comparisons. To visualize the results, we generated boxplots for each alignment method, overlaying individual data points with jitter to illustrate the spread. Significant differences between groups were annotated using asterisk notation to indicate the level of statistical significance (p < 0.05 *, p < 0.01 **, p < 0.001 ***).

#### Gene expression similarity between aligned spatial locations

For each matched pair of spots or bins, we extracted gene expression vectors from the molecular data for both sections and evaluated their similarity using cosine similarity. We first identified highly variable genes of the reference section using log-transformed expression values in Scanpy (sc.pp.highly_variable_genes)^40^. Cosine similarity was computed between the highly-variable gene expression vectors for each spot pair. This analysis was performed across five different alignment methods (COAST, Unaligned, Rigid alignment, MOSCOT, CAST). Results were visualized using a kernel density estimation plot (*seaborn.kdeplot*).

#### Neighborhood Preservation Score

To assess tissue distortion introduced by the alignment process, we quantified neighborhood preservation by comparing spatial neighborhoods before and after alignment. For the query section (pre-alignment and post-alignment), we identified the nearest neighbors of each spot using a k-nearest neighbors approach with *k* set to 24 for Visium data and *k* set to 48 for Stereo-seq. Specifically, for each spot, the *k* closest neighboring spots were determined based on Euclidean distance in the two-dimensional coordinate space, using the KD-tree algorithm. Spatial coordinates prior to alignment were obtained from the original dataset, while coordinates after alignment were taken from the dataset aligned with COAST. The distribution of overlapping neighbors across all spots was visualized using a bar plot showing the frequency of spots by the count of overlapping neighbors.

### Ischemic-reperfusion injury (IRI) study

The experiment was carried out as in Rietjens et al. 2025 ^33^ and Wang et al. 2022 ^3^. Experiments consisted of three 12-week-old male constitutional renin reporter (B6.Ren1cCre/TdTomato/J) mice ^41^. The renal vasculature was clamped for approximately 18 minutes. Clamp removal allowed blood flow to resume (reperfusion) and the abdomen was closed. At day 14 post-surgery, mice were sacrificed by perfusion with cold PBS-heparin (5 UI/mL) via the left ventricle for 6 minutes at a controlled pressure of 150 mmHg before removing the kidney. MALDI-MSI and Stereo-seq sections were taken 10 μm apart. Animal experiments were approved by the Ethical Committee on Animal Care and Experimentation of the Leiden University Medical Center (permit no. AVD1160020171145). The Stereo-seq data generation is described in detail in Rietjens et al. 2025 ^33^.

#### MALDI-MSI Tissue Preparation and Matrix Deposition

Tissue slices were embedded in 10% gelatin and cryosectioned into 10 µm thick sections using a cryostat at -20 °C. The sections were thaw-mounted onto indium-tin-oxide (ITO)-coated glass slides (VisionTek Systems Ltd., Chester, UK). Mounted sections were placed in a vacuum freeze-dryer for 15 minutes prior to matrix application. After drying, N-(1-naphthyl) ethylenediamine dihydrochloride (NEDC) (Sigma-Aldrich, UK) MALDI-matrix solution of 7 mg/mL in methanol/acetonitrile/deionized water (70, 25, 5 %v/v/v) was applied using a HTX M3+ Sprayer (HTX Technologies, USA). The setting was as below: temperature, 60 °C; number of passes, 20 layers; flow rate, 80 μL/min; velocity, 2000 mm/min; track spacing, 3 mm; gas flow rate 10 psi, and drying time in between passes, 30 s.

#### MALDI-MSI measurement and data analysis

MALDI-TOF/TOF-MSI was performed using a RapifleX MALDI-TOF/TOF system (Bruker Daltonics GmbH, Bremen, Germany). Negative ion-mode mass spectra were acquired at a pixel size of 10 × 10 µm^2^ over a mass range from m/z 80-1000. Prior to analysis the instrument was externally calibrated using red phosphorus. Spectra were acquired with 50 laser shots per pixel at a laser repetition rate of 10 kHz. Data acquisition was performed using flexControl (Version 4.0, Bruker Daltonics) and visualizations were obtained from flexImaging 5.0 (Bruker Daltonics). MSI data were exported and processed in SCiLS Lab 2024b pro (SCiLS, Bruker Daltonics) with baseline correction using convolution algorithm. All MALDI-TOF-MSI data were normalized to the Root Mean Square (RMS). High-molecular-weight features (m/z > 400, predominantly phospholipids) were picked at signal-to-noise-ratio > 3 on the average spectrum, and matrix peaks were excluded from the m/z feature list. The m/z values from MALDI-TOF were imported into the Human Metabolome Database (https://hmdb.ca/) after recalibration in mMass and annotated for metabolites with an error < ±20 ppm. Peak intensities of the selected features were exported for all the measured pixels from SCiLS Lab, which were used for the following analysis.

#### Stereo-seq data preprocessing

The design, tissue segmentation, and preprocessing of the Stereo-seq experiments, including cell-type annotation, were performed as described in Rietjens et al. ^33^. Importantly, the spatial coordinates, and the corresponding count matrix, were binned using 40 as bin size (20 µm x 20µm resolution), resulting in new spot identifiers. The final Stereo-seq datasets contained 48,347, 40,918, and 40,494 spatial spots for IRI1, IRI2, and IRI3, respectively. The image associated with the Stereo-seq experiment comprised all three tissue sections and was segmented into individual regions for downstream analysis, yielding image sizes of 12,659 x 11,336 pixels for IRI1, 9,710 x 15,063 pixels for IRI2, and 10,444 x 12,561 pixels for IRI3. The three replicates were corrected for batch effect using rPCA.

#### Stereo-seq ssDNA image preprocessing and segmentation

In Figure 4b, we observe that the ssDNA tissue images from Stereo-seq exhibit artifacts (a grid-like pattern of squares resulting from the imaging process). To address this, we performed cell segmentation on the Stereo-seq image and applied a binary mask, where detected cells are marked in white and all other areas in black (Supplementary Fig. S6). More in detail, whole-slide mouse kidney tissue DAPI-stained images were processed using a custom nuclei segmentation pipeline implemented in Python 3. The pipeline employs the nuclei model of Cellpose3 ^42^, combined with a tile-based strategy, in which the image is split into overlapping tiles, for scalable and adaptive mask extraction. In the preprocessing stage non-tissue background pixels were manually zeroed using Fiji ^43^. The images are enhanced using contrast-limited adaptive histogram equalization (CLAHE) ^44^ to improve nucleus-background separability and then split into overlapping square tiles. Tile size and overlap are determined by the parameters “tile_side_length” and “tile_overlap,” set to 512 and 0.2, respectively. Tiling enables memory-efficient processing of arbitrarily large images and supports adaptive parameterization. In this study, we utilized this adaptability for Cellpose3’s diameter parameter, which is estimated from the median equivalent diameter of objects initially segmented via flow fields and conveys to the model what average nuclei diameters to expect. To ensure that nuclei are not split across tile borders, the overlap must be large enough to fully capture even the largest nuclei but not so large as to cause the borders of non-adjacent tiles to approach each other. Consequently, we chose a tile size of 512 pixels to balance accuracy, memory load, and estimation stability in our systems and images.

Tiles are processed in batches through the Cellpose3 “nuclei” model. To increase sensitivity, the cell probability threshold was lowered from the default 0 to -14, CLAHE contrast was intensified (clip limit = 5.0, grid size = 32x32), and the flow threshold was raised from the default 0.4 to 0.9, requiring more coherent flow fields for a nucleus to be accepted. This parameter combination implements a sensitivity-specificity tradeoff, with the flow threshold acting as the principal specificity regulator. Parameter settings were determined by evaluating performance on annotated image subsets, using Intersection-over-Union (IoU) as the criterion for segmentation quality. For adaptive diameter estimation to function, diameter was set to None and resample to True. All other Cellpose3 parameters remained at their default values.

To reconstruct the full mask set for the original image, a custom tile-merging algorithm was applied. For each overlapping tile pair, the following four steps were taken sequentially:

1. Priority Tile Selection: The tile with the greater number of detected nuclei is designated as the priority tile to minimize inclusion of edge-adjacent masks, which are more likely to be affected by border artifacts.
2. Priority Tile Border Mask Removal: The location of the priority tile’s edge within the non-priority tile is identified. All priority tile masks intersecting this boundary are removed to avoid including potentially distorted nuclei.
3. Preservation of Boundary Non-Priority Masks: Non-priority tile masks intersecting the corresponding boundary are marked down to be preserved.
4. Removal of Redundant Non-Priority Masks: All non-priority tile masks within the overlapping region that were not marked for preservation are discarded.

This process ensures that masks in the overlapping area originate only from the priority tile, except along its border, where non-priority masks are used to avoid edge artifacts. The method effectively prevents merging into oversized masks, duplication, and artifacts from the tile creation while being computationally efficient, even on CPU systems.

Finally, we used an optional filtering script to evaluate each mask based on its circularity, eccentricity, size, solidity, hole fraction, and aspect ratio. Masks with values outside user-defined thresholds were removed from the final output. We used the following threshold: min_pixels=20, max_pixels=900, min_circ=0.56, max_circ=1.00, min_sol=0.765, max_sol=1.00, min_ecc=0.00, max_ecc=0.975, min_ar=0.50, max_ar=3.20, min_hole=0.00, max_hole=0.001.

#### MALDI-MSI preprocessing

The final datasets included 240,255 spots for IRI1, 161,596 for IRI2, and 165,870 for IRI3, at 10 µm x 10µm resolution. Deisotoping was performed manually in SCiLS software. After deisotoping, a total of 301 lipid and 285 metabolite features were retained for all three tissues. The corresponding image dimensions were 4,352 x 5,120 pixels for IRI1, 4,352 x 4,864 pixels for IRI2, and 3,840 x 4,352 pixels for IRI3. The three replicates were corrected for batch effect with combat (implemented in scanpy.pp.combat). ^45^

#### Stereo-seq cell type - m/z marker analysis

Lipid cell-type markers were found in literature. More in detail, from Farrow et al. 2025 ^34^ we selected the top 10 markers per cell type (glomeruli, TAL, PT, Distal, Collecting duct) that differed no more than +/- 0.1 m/z from our detected features. From Wang et al. 2022 ^27^ we retained all markers within a 0.1 m/z tolerance. Average lipid marker expression per cell type was visualized using custom heatmaps. For each feature, expression values were extracted from the MSI data, averaged across cell types (cell-type annotation obtained as described in Rietjens et al. 2025 ^33^), and standardized by z-score transformation.

#### Metabolites and lipids differential expression analysis

Differential gene expression analysis was performed using the Wilcoxon rank-sum test implemented in Scanpy (sc.tl.rank_genes_groups), grouping cells by transcriptomics-derived cell type annotations. The top 10 marker genes per cell type were identified based on ranked test statistics, using root mean squared-normalized expression values.

## Data Availability

10X Mouse Brain Sagittal Posterior (Visium). Section 1: https://www.10xgenomics.com/datasets/mouse-brain-serial-section-1-sagittal-posterior-1-standard-1-0-0. Section 2: https://www.10xgenomics.com/datasets/mouse-brain-serial-section-2-sagittal-posterior-1-standard

10X Mouse Brain Coronal (Visium). Section 1: https://www.10xgenomics.com/datasets/mouse-brain-coronal-section-1-ffpe-2-standard Section 2: https://www.10xgenomics.com/datasets/mouse-brain-coronal-section-2-ffpe-2-standard

10X Mouse Brain Sagittal Anterior (Visium). Section 1: https://www.10xgenomics.com/datasets/mouse brain-serial-section-1-sagittal-anterior-1-standard-1-1-0. Section 2: https://www.10xgenomics.com/datasets/mouse-brain-serial-section-2-sagittal-anterior-1-standard

MOSTA Mouse Embryo: https://db.cngb.org/stomics/mosta/download/

Stereo-seq dataset: GSE298899

Stereo-seq images: https://doi.org/10.6084/m9.figshare.30720194.v1

MALDI-MSI dataset: https://zenodo.org/records/17722895

Pretrained ssDINO Vision Transformer: https://github.com/facebookresearch/dino (Architecture: ViT-S/16)

## Code Availability

Code for COAST and figure reproducibility is available at https://github.com/mahfouzlab/COAST.

## Acknowledgements

We acknowledge Manon Zuurmond (Department of Internal Medicine, LUMC, Leiden, the Netherlands) for making the illustrations for this article. The Novo Nordisk Foundation Center for Stem Cell Medicine (reNEW) is supported by Novo Nordisk Foundation grants (NNF21CC0073729). S.D. is supported by funding from Agence Nationale de la Recherche (ANR24-CPJ1-0181-01). T.R. is supported by the European Union through ERC grant (SPARK 101140863). Views and opinions expressed are however those of the author(s) only and do not necessarily reflect those of the European Union or the European Research Council Executive Agency. Neither the European Union nor the granting authority can be held responsible for them.

