## Supplementary Figures S1-S7 for "Image-guided alignment of consecutive multi-modal tissue slides"

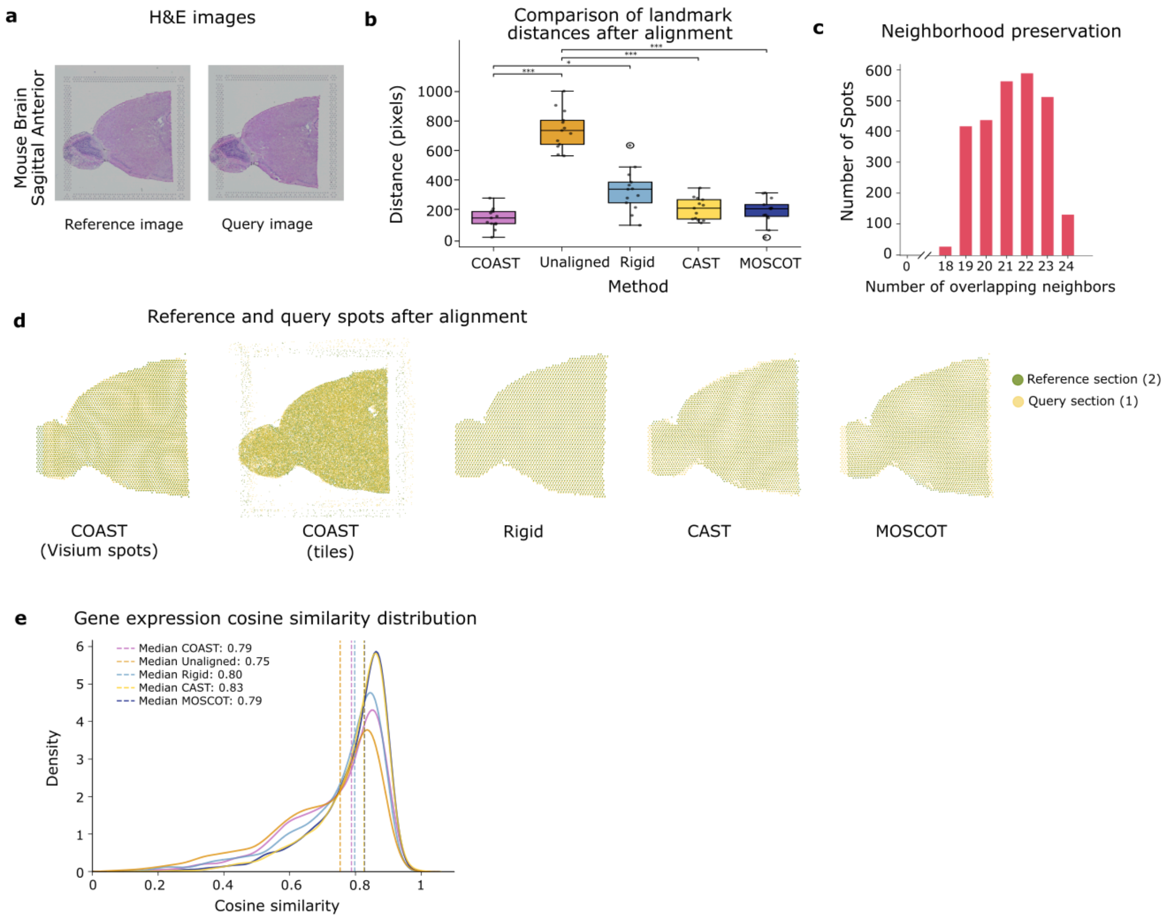


**Supplementary Figure S1**: **Results of the alignment with COAST on 10X Mouse Brain Sagittal Anterior (Visium)**. **a**: H&E images of the two consecutive mouse brain sections. **b**: Boxplot representing the distances between landmarks after registration via COAST, Rigid Alignment, Unaligned, CAST and MOSCOT. **c**: Neighborhood preservation of the third order neighbourhood (24 spots) in the query section before and after alignment. **d**: Spatial distribution of reference and query spots after alignment with COAST and benchmarking tools. **e**: Cosine similarity of gene expression of matched spots with COAST, rigid alignment, CAST and MOSCOT (raw counts, 3000 highly variable genes).

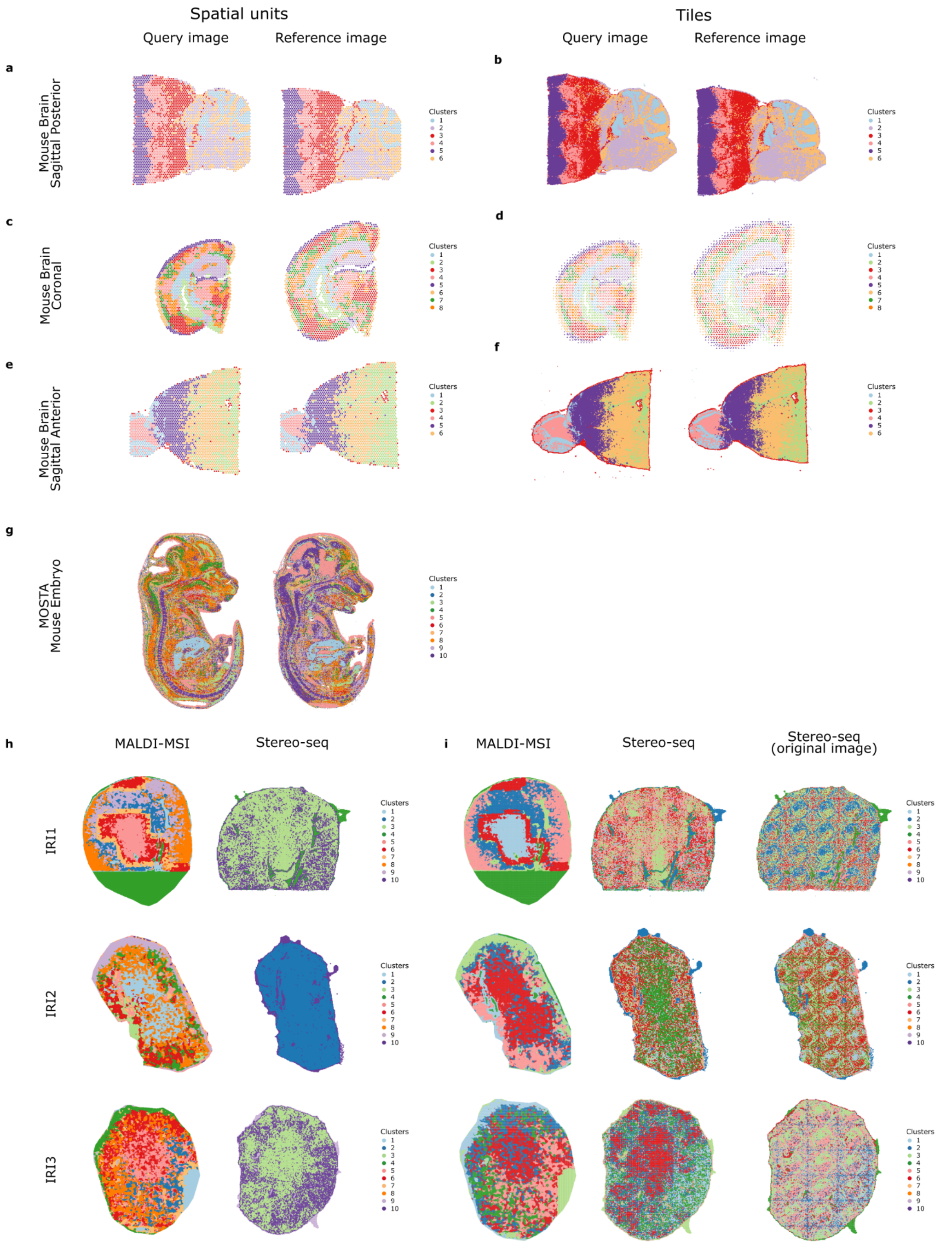


**Supplementary Figure S2: K-means clustering applied on the vision transformer embedding**. **a**: Clustering on spatial transcriptomics spots from 10X Mouse Brain Sagittal Posterior dataset. **b**: Clustering on tiles from 10X Mouse Brain Sagittal Posterior dataset. **c**: Clustering on spatial transcriptomics spots from 10X Mouse Brain Coronal dataset. **d**: Clustering on tiles from 10X Mouse Brain Coronal dataset. **e**: Clustering on spatial transcriptomics spots from 10X Mouse Brain Sagittal Anterior dataset. **f**: Clustering on tiles from 10X Mouse Brain Sagittal Anterior dataset. **g**: Clustering on spatial transcriptomics units from MOSTA Mouse Embryo dataset. **h**: Joint clustering on tiles from IRI mouse kidney dataset (tissue IRI1, IRI2, IRI3). Left: MALDI-MSI embedding, right: Stereo-seq (cell-segmented image) embedding. **i**: Individual clustering on mouse kidney dataset (tissue IRI1, IRI2, IRI3). Left: MALDI-MSI embedding, center: Stereo-seq (cell-segmented image) embedding, right: Stereo-seq (original image) embedding.

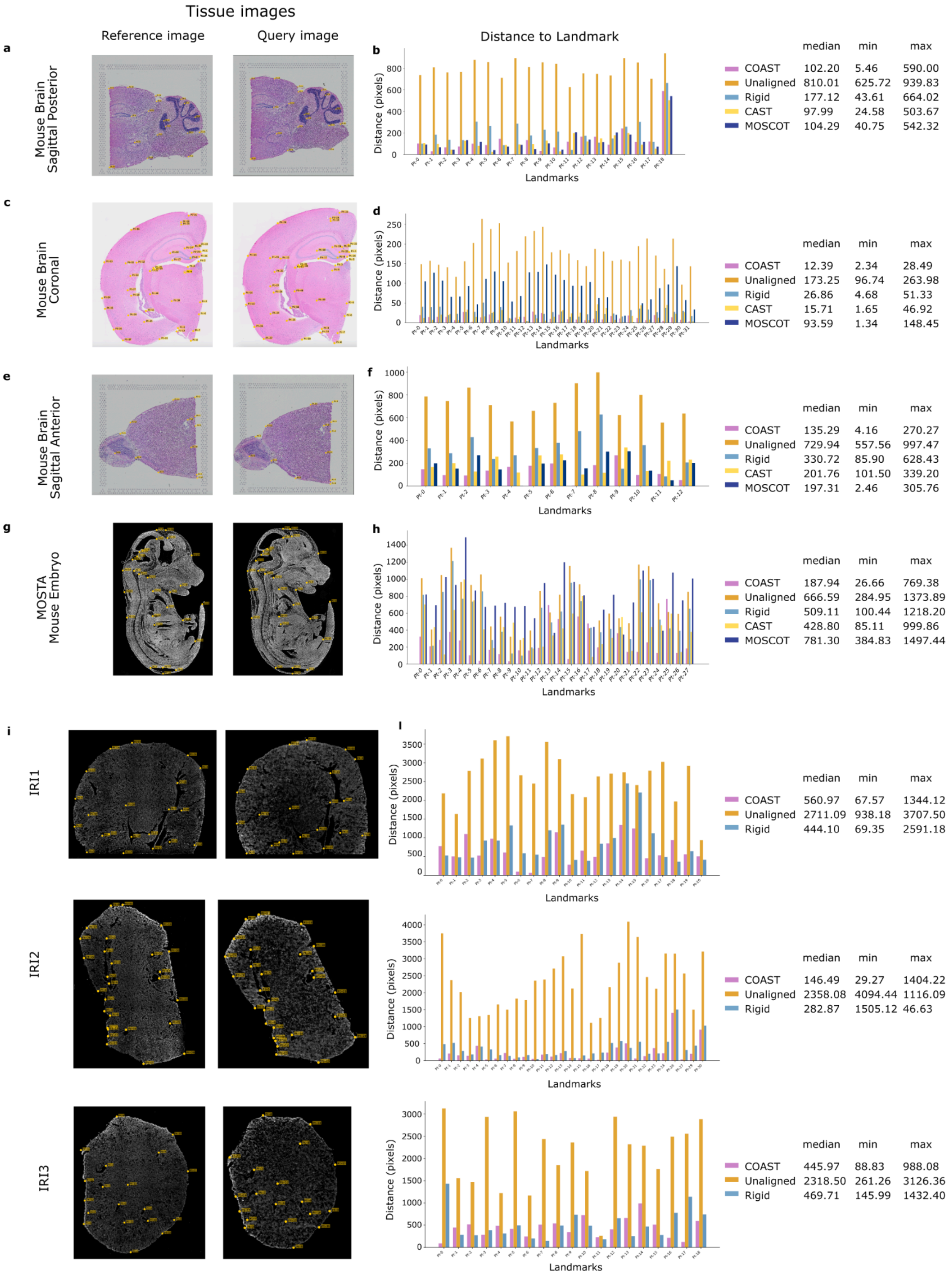


**Supplementary Figure S3**. **Landmarks position on tissue and landmark distances after alignment**. Left column: Landmark locations on tissues, query: left, reference: right. **Central column**: Barplot of individual landmark euclidean distances for COAST, Rigid alignment, Unaligned, CAST and MOSCOT. **Right column**: statistics (median, min, max) on landmark distances. **a,b**: Mouse Brain Sagittal Posterior, **c,d**: Mouse Brain Coronal, **e,f**: Mouse Brain Sagittal Anterior, **g,h**: MOSTA Mouse Embryo, **i,l**: IRI mouse kidney dataset (tissues IRI1, IRI2, IRI3)

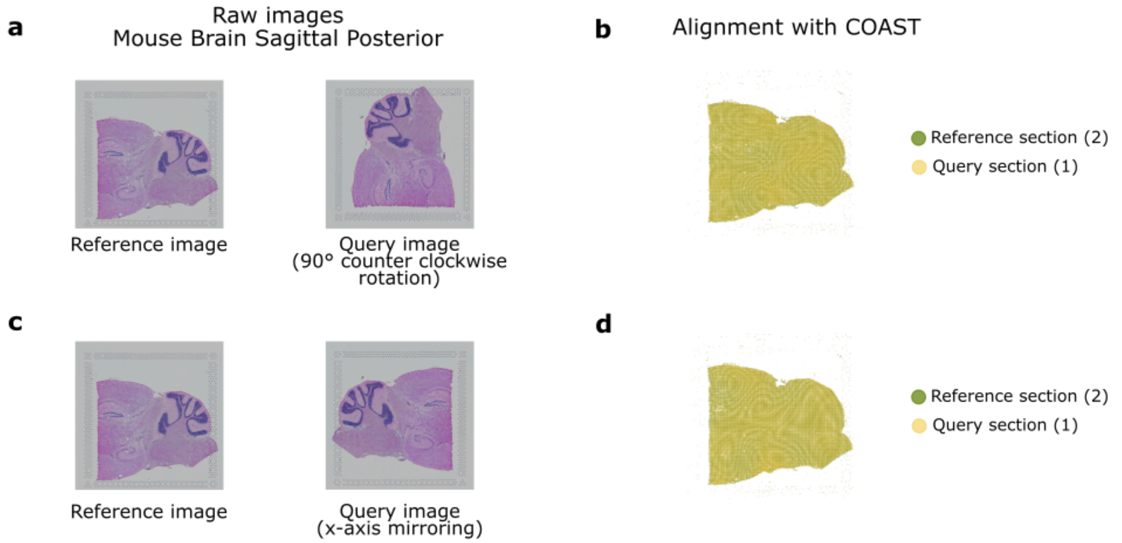


**Supplementary Figure S4**: **Results of the alignment with COAST on 10X Mouse Brain Sagittal Posterior. a**: H&E input images, query has been rotated 90° counter clockwise. **b**: Query and reference sections after alignment with COAST. Left: tiles, right: spatial transcriptomics spots. **c**: H&E input images, query has been mirrored around the x-axis . **d**: Query and reference sections after alignment with COAST.

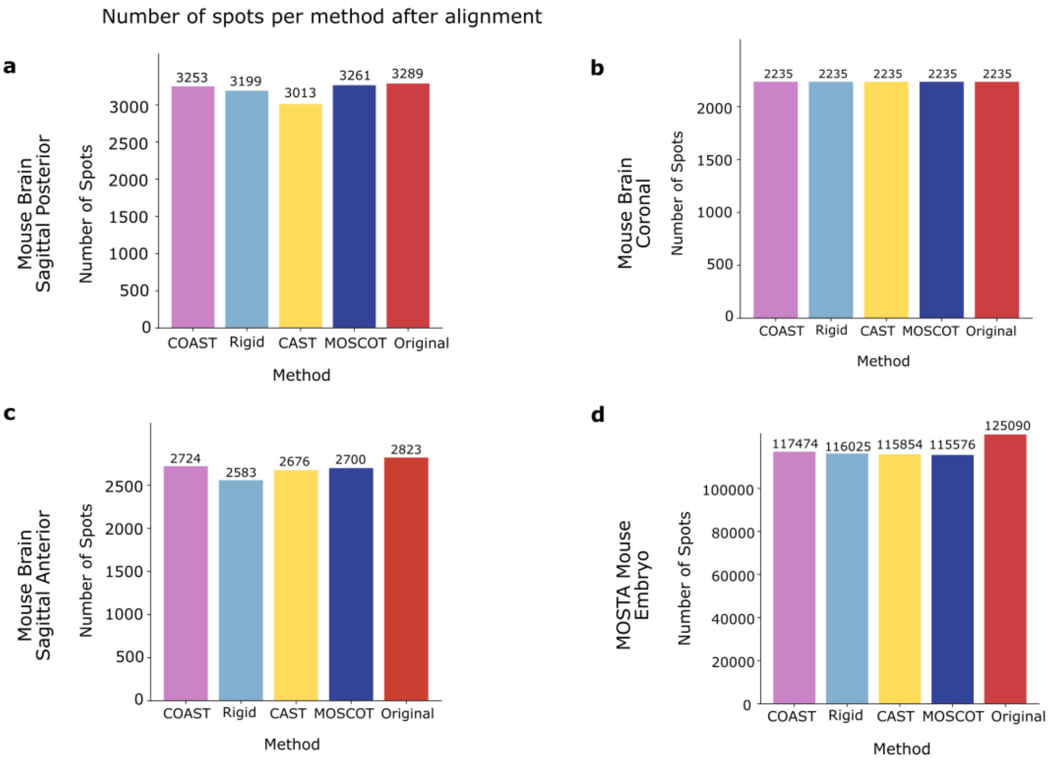


**Supplementary Figure S5. Number of spots in the matched dataset after alignment. a**: 10X Mouse Brain Sagittal Posterior. **b**: 10X Mouse Brain Coronal. **c**: 10X Mouse Brain Sagittal Anterior. **d**: Stereo-seq MOSTA Mouse Embryo.

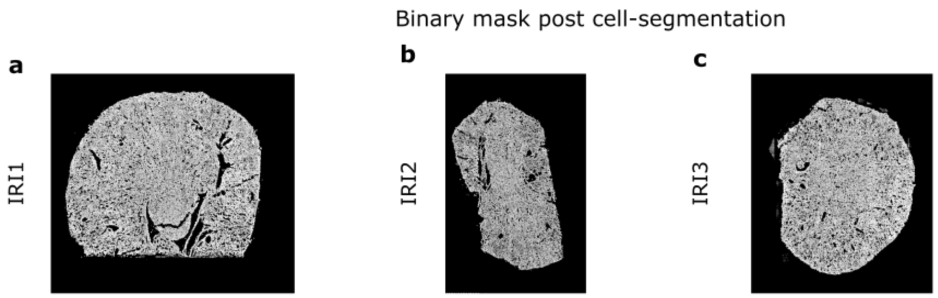


**Supplementary Figure S6**. **Binary mask post-cell segmentation overlaid on the Stereo-seq ssDNA images** for IRI1 (**a**), IRI2 (**b**), IRI3 (**c**).

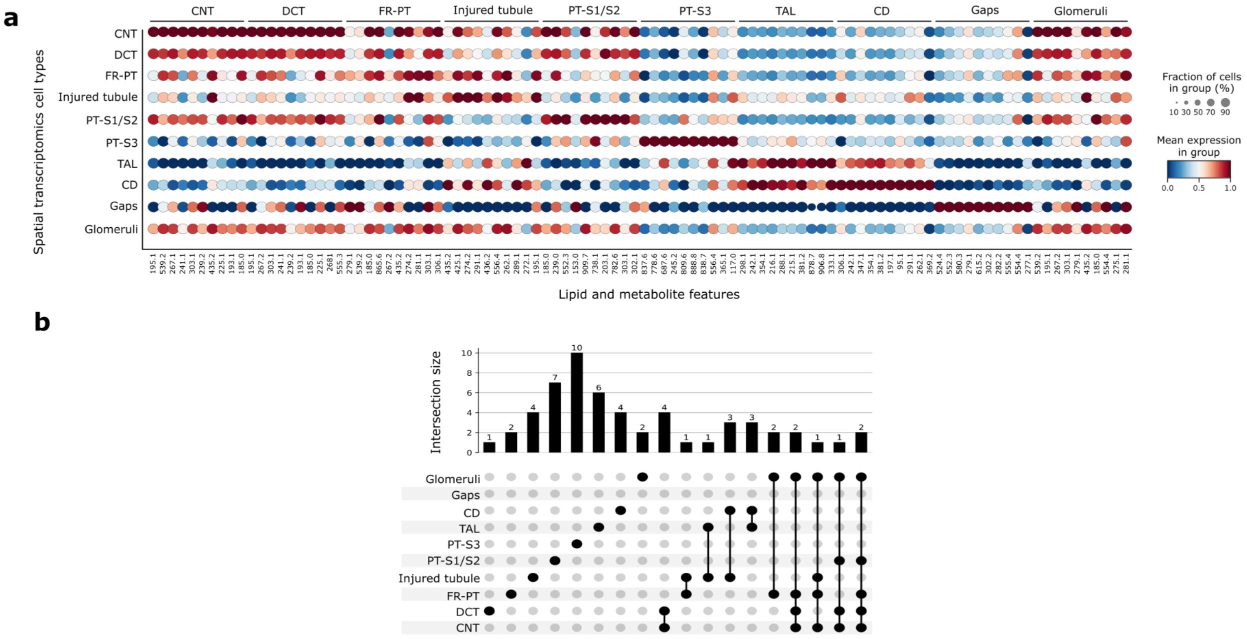


**Supplementary Figure S7. a**: dot plot of top 10 differentially expressed lipids and metabolites per cell type. **b**: Shared top differentially expressed markers across renal cell types. UpSet plot showing intersections among the top 10 differentially expressed features per cell type. Each bar indicates the number of markers shared between one or more groups.
